# Distinguishing signal from autofluorescence in cryogenic correlated light and electron microscopy of mammalian cells

**DOI:** 10.1101/120642

**Authors:** Stephen D. Carter, Shrawan K. Mageswaran, Zachary J. Farino, João I. Mamede, Catherine M. Oikonomou, Thomas J. Hope, Zachary Freyberg, Grant J. Jensen

## Abstract

Cryogenic correlated light and electron microscopy (cryo-CLEM) is a valuable tool for studying biological processes *in situ*. In cryo-CLEM, a target protein of interest is tagged with a fluorophore and the location of the corresponding fluorescent signal is used to identify the structure in low-contrast but feature-rich cryo-EM images. To date, cryo-CLEM studies of mammalian cells have relied on very bright organic dyes or fluorescent protein tags concentrated in virus particles. Here we describe a method to expand the application of cryo-CLEM to cells harboring genetically-encoded fluorescent proteins. We discovered that a variety of mammalian cells exhibit strong punctate autofluorescence when imaged under cryogenic conditions (80K). Compared to fluorescent protein tags, these sources of autofluorescence exhibit a broader spectrum of fluorescence, which we exploited to develop a simple, robust approach to discriminate between the two. We validate this method in INS-1 E cells using a mitochondrial marker, and apply it to study the ultrastructural variability of secretory granules in a near-native state within intact INS-1E pancreatic cells by high-resolution 3D electron cryotomography.

## Introduction

Electron microscopy (EM) is an essential tool in the study of cell ultrastructure, with resolving power several orders of magnitude greater than that of light microscopy (LM). Frequently, however, it is difficult to identify objects of interest in EM images when their ultrastructure is unknown. In conventional “thin-section” transmission electron microscopy (TEM), this challenge was addressed by the development of immuno-gold labeling (Faulk and Taylor 1971). Although this method allows direct labeling and visualization of specific targets within the cell, the fixation, dehydration and/or resin embedding steps can result in poor cell and antigen preservation and accompanying loss of information.

Alternatively, in correlated light and electron microscopy (CLEM), each target can be specifically labelled with a genetically-encoded fluorescent protein (Briegel, Chen et al. 2010), located first by fluorescence light microscopy, and then imaged at higher magnification by electron microscopy. CLEM can be done either at room- or cryogenic-temperatures (“cryo-CLEM”). Like immunoEM, room-temperature CLEM also requires chemically fixed and dehydrated cells, which can distort or obscure important structural features (Afzelius and Maunsbach 2004, Lucic, Rigort et al. 2013), but it has nevertheless allowed the visualization of numerous bacterial and mammalian cellular events that would otherwise have been challenging or impossible to capture (Voloshin Ia, Suslov Ie et al. 2000, Grabenbauer, Geerts et al. 2005, Darcy, Staras et al. 2006, Kapoor, Lampson et al. 2006, Muller-Reichert, Srayko et al. 2007, Kukulski, Schorb et al. 2011, Kukulski, Schorb et al. 2012, Redemann and Muller-Reichert 2013, Avinoam, Schorb et al. 2015, Bertipaglia, Schneider et al. 2016, Schorb, Gaechter et al. 2016).

In cryo-CLEM samples are preserved in a near-native, “frozen-hydrated” state. To visualize fluorescence inside frozen-hydrated cells, cryogenic LM (cryo-LM) stages are used (Briegel,Chen et al. 2010, Schlimpert, Klein et al. 2012, Schellenberger, Kaufmann et al. 2014, Schorb and Briggs 2014, Bertipaglia, Schneider et al. 2016, Schorb, Gaechter et al. 2016). Unfortunately, because the sample has to be kept frozen, long-working-distance air objective lenses with low numerical apertures are used instead of oil-immersion lenses. To increase the resolution of the light microscopy, several “super-resolution” cryo-CLEM studies have also now been performed (Chang, Chen et al. 2014, Liu, Xue et al. 2015).

To date, cryo-CLEM studies of mammalian cells have used either very bright organic fluorescent dyes or fluorescent proteins concentrated in viruses (Jun, Ke et al. 2011, Schellenberger, Kaufmann et al. 2014, Bykov, Cortese et al. 2016). Organic dyes are very bright, but it can be challenging to attach them to specific proteins inside cells. Genetically-encoded fluorescent proteins, on the other hand, can be easily fused with most proteins and offer a richer repertoire of colors, but they are much less bright, so their signals can be more difficult to distinguish from cellular autofluorescence. Here we describe a method to distinguish signal arising from genetically-encoded fluorescent proteins from endogenous autofluorescence in mammalian cells under cryogenic conditions. We validate the approach with mitochondria and then demonstrate how the method allows secretory granules to be identified and structurally characterized within intact pancreatic cells with unprecedented resolution.

## Results

### INS-1E cells exhibit bright, punctate autofluorescence at ∼80K

Initially, we set out to image the secretory pathway of the pancreatic beta cell-derived INS-1E line, which has long been used as a model system (Merglen, Theander et al. 2004, Farino, Morgenstern et al. 2016). The secretory machinery of these cells has been well studied by both LM and conventional EM (Rubi, Ljubicic et al. 2005, Giordano, Brigatti et al. 2008), but we wanted to advance that by imaging cells in a near-native, frozen-hydrated state in 3-D by using electron cryotomography (ECT) (Oikonomou and Jensen 2016). We were particularly interested in dense core secretory granules (DCSGs), which are central to the efficient secretion of hormones including insulin (Kim, Tao-Cheng et al. 2001). To identify DCSGs in cryotomograms, we tagged chromogranin A (CgA), a granin protein widely used as a marker for DCSGs given its almost exclusive localization to this intracellular compartment (Huh, Bahk et al. 2005).

We transfected INS-1E cells with CgA C-terminally tagged with GFP and grew cells to around 30-40% confluency on EM finder grids. To facilitate image correlation we added 500 nm blue fluorospheres visible in both cryo-LM and cryo-EM modalities to the samples before plunge-freezing [as in (Chang, Chen et al. 2014, Schellenberger, Kaufmann et al. 2014, Schorb and Briggs 2014, Liu, Xue et al. 2015, Bykov, Cortese et al. 2016)]. We first performed cryo-LM to identify targets of interest in thin ice at the periphery of cells. To accommodate the relatively dim signal of CgA-GFP (due either to the low temperature or the low NA of our long working-distance air objective (NA 0.7), or both), we used exposure times of up to 2 seconds and applied a 2D real-time deconvolution algorithm. We observed a punctate cytosolic distribution (Figure 1A) consistent with CgA’s expected intracellular localization to secretory granules (Huh, Bahk et al. 2005).

As a negative control, we also imaged untransfected INS-1E cells by cryo-LM (Figure 1b). Unlike room-temperature images of untransfected cells, which display little autofluorescence (Figure 1c-d), to our surprise unlabelled INS-1E cells exhibited clear puncta at ∼80K (cooled by liquid nitrogen). To determine whether the autofluorescence observed at 80K was broad-spectrum, as is typical at room-temperature (Billinton and Knight 2001), we also imaged untransfected and transfected cells using an mCherry filter. In transfected cells there were two populations of puncta: one emitting both green and red fluorescence, and one emitting primarily green fluorescence (Figure 1a). In untransfected cells, all the puncta emitted both green and red fluorescence (Figure 1b), indicating that the autofluorescence in the sample was broad spectrum.

**Figure 1.**
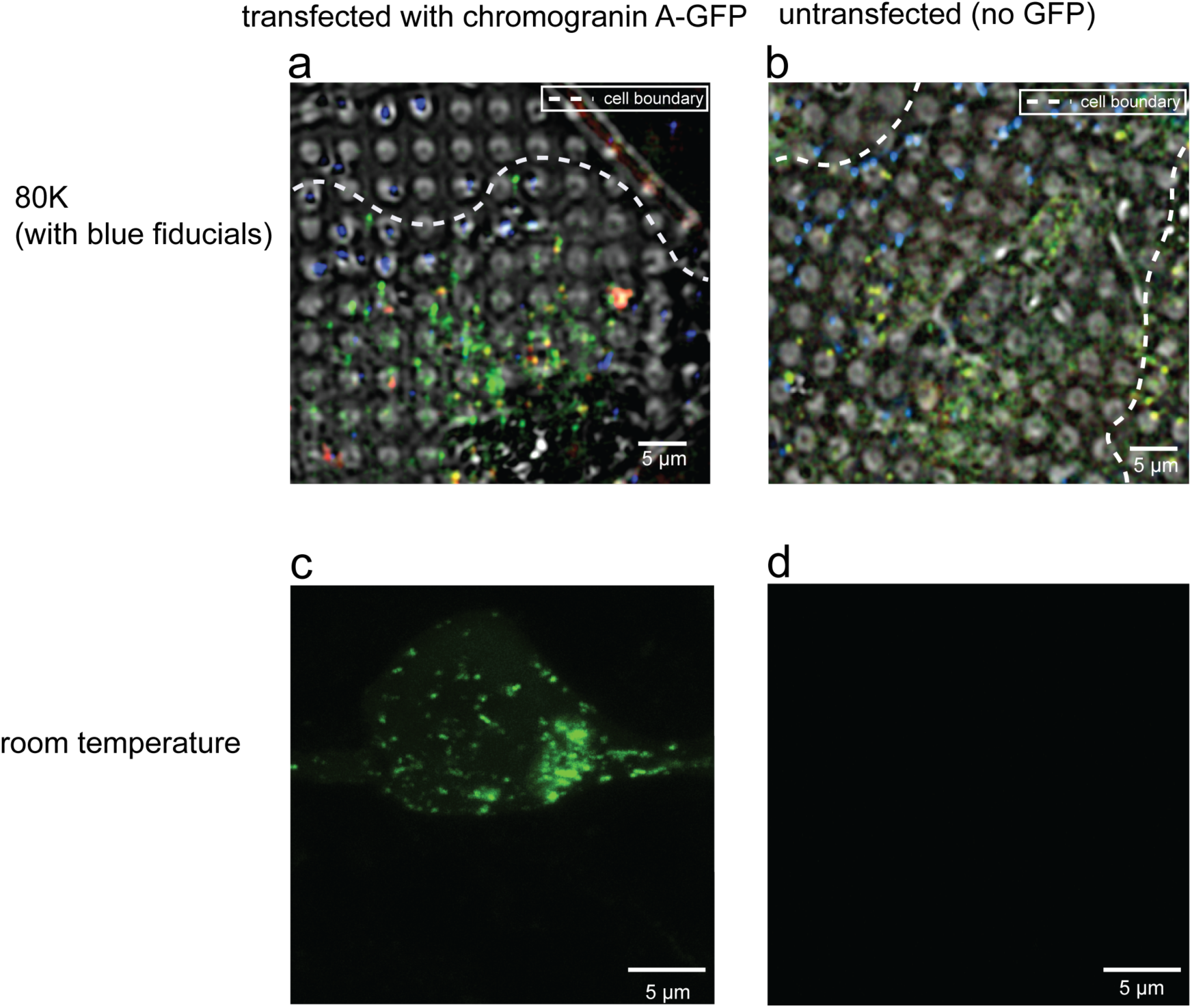
INS-1E cells exhibit strong autofluorescence at 80K. **(a,b)** Cryo-LM images (composite of bright field and epifluorescence in FITC, mCherry and DAPI channels) of INS-1E cells transfected with CgA-GFP (A) or untransfected (B). **(c,d)** Room-temperature light microscopy images of epifluorescence in FITC channel of INS-1E cells transfected with CgA-GFP (C) or untransfected (D).

### Other mammalian cells also exhibit bright autofluorescence at ∼80K

Next we checked if bright autofluorescence at 80K was unique to INS-1E cells. We applied the same imaging technique to three other untransfected cell lines: rhesus macaque fibroblasts, HeLa cells and human primary adipocytes. All three cell lines exhibited numerous puncta distributed throughout the cell volume with broad fluorescence spectra ranging from green (FITC) to red (mCherry) (Figure 2). To identify the source of the observed autofluorescence, we imaged several puncta at high-resolution by ECT. All of the autofluorescent puncta in rhesus macaque fibroblasts (Figure 3a) and HeLa cells (data not shown) correlated to multimembranous structures, some resembling multilamellar bodies (MLBs) (Hariri, Millane et al. 2000, Lajoie, Guay et al. 2005). Similarly, many autofluorescent puncta in untransfected INS-1E cells also correlated to multimembranous structures in cryotomograms (Figure 3b, panel 1-3). Other autofluorescent puncta in INS-1E cells correlated to membrane-enclosed crystals (Figure 3b, panel 4).

**Figure 2.**
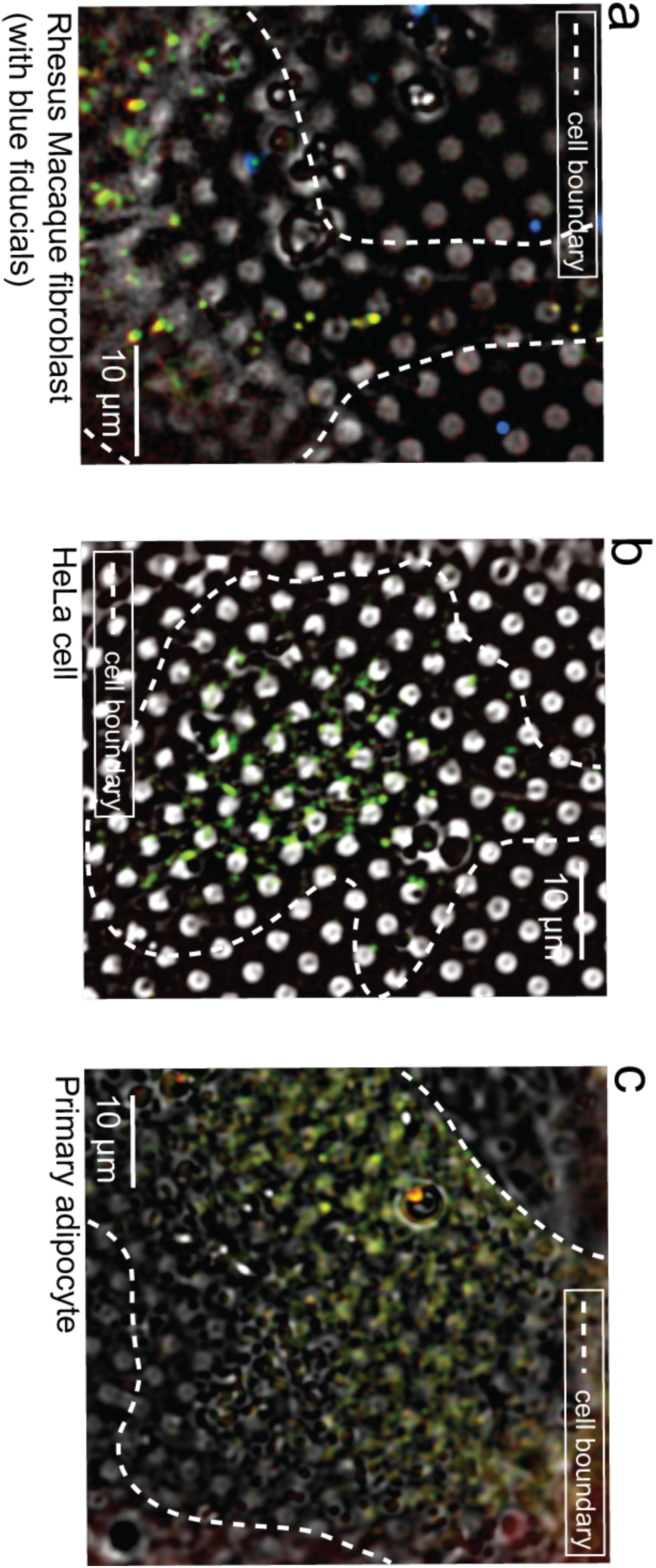
Autofluorescence at 80K is a general feature of mammalian cells. Cryo-LM images (bright field and epifluorescence in FITC, mCherry and DAPI channels) of **(a)** rhesus macaque fibroblasts, **(b)** HeLa cells and **(c)** primary adipocytes.

**Figure 3.**
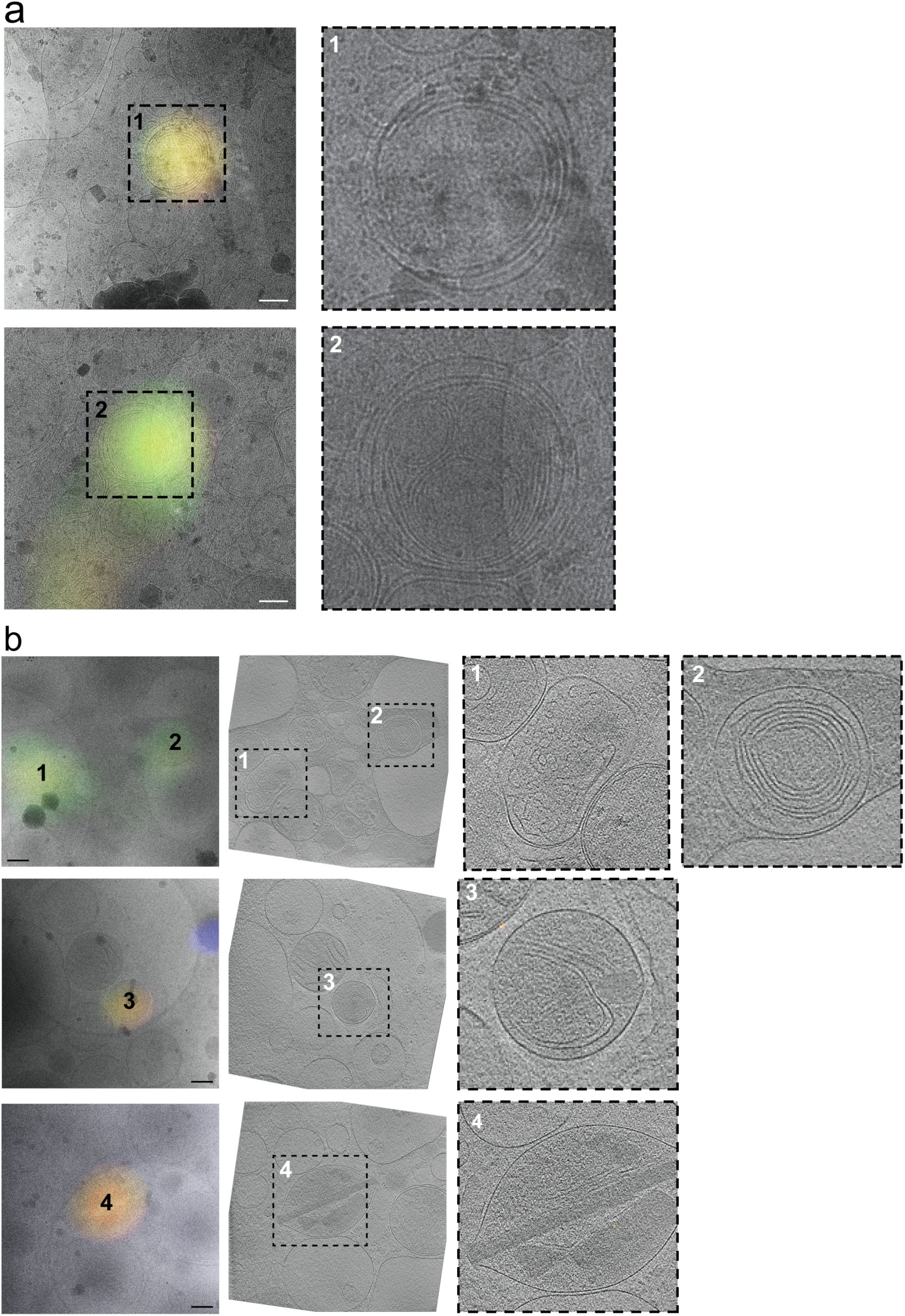
Cryo-CLEM reveals sources of autofluorescence in untransfected rhesus macaque fibroblasts (a) and INS-1E cells (b). Left panels show epifluorescence images overlaid on high-magnification cryo-EM projection images. Middle panel in (b) shows the corresponding tomographic slice. Right panels show zoomed-in views of the boxed areas in corresponding panels at left. Numbers indicate corresponding locations. Scale bars = 200 nm.

### Distinguishing fluorescent protein tags from autofluorescence

At room temperature, the confounding effect of cellular autofluorescence can sometimes be mitigated by photobleaching; background fluorescence sources tend to bleach faster than fluorescent proteins (Billinton and Knight 2001). Unfortunately, the autofluorescence we observed at 80K did not bleach away quickly (data not shown), as observed previously for exogenous fluorophores (Schwartz, Sarbash et al. 2007). Instead, we tried to identify autofluorescence by its broader emission spectrum, as has been done previously at room temperature (Szollosi, Lockett et al. 1995, Mansfield, Gossage et al. 2005).

To test the method, we chose a “positive control” that could be unambiguously identified in cryotomograms with or without a fluorescent tag. Mitochondria proved to be a good choice due to their well-defined ultrastructure including easily-recognizable cristae (Rabl, Soubannier et al. 2009, Zick, Rabl et al. 2009, Davies, Daum et al. 2014). We transfected INS-1E cells with the mitochondrial marker Mito-DsRed2, which is targeted to the space between the inner and outer mitochondrial membrane by the signal sequence of cytochrome C oxidase. We again imaged both untransfected and transfected cells by cryo-CLEM, and measured the fluorescence of a large number of puncta in both the green (FITC) and red (mCherry) channels. While both samples exhibited puncta emitting in both the mCherry and FITC channels, there was a population only in the transfected cells that exhibited high intensity in the mCherry channel and low intensity in the FITC channel (Figure 4a, top left corner), indicative of specific Mito-DsRed2 fluorescence.

To identify structures associated with fluorescence in these cells, we imaged 32 randomly chosen puncta in transfected cells by ECT. Many of the puncta correlated to mitochondria, and the centroid of fluorescence fell within the boundaries of the organelle, validating the accuracy of our correlation procedure (Figure 4b, targets 1-7). Others correlated to multimembranous structures (Figure 4b, targets 8-11). As expected, puncta correlating to mitochondria exhibited a greater ratio of red fluorescence to green fluorescence, and were therefore positioned near the upper left corner of the fluorescence plot (Figure 4a). The multimembranous structures corresponded to puncta near the opposite (lower right) corner. Thus, to distinguish fluorescent protein tags from autofluorescence, one can record fluorescence images of both tagged and untagged frozen cells and plot the fluorescence in two channels. Puncta in the overlap region near the middle of the plot may be from either fluorescent protein tags or autofluorescence, and cannot be distinguished with great confidence. Puncta in the extreme corner towards the pure color of the fluorescent protein tag where autofluorescent puncta are not seen (triangle in the upper left corner of Figure 4a), however, are very likely to contain the tag.

**Figure 4.**
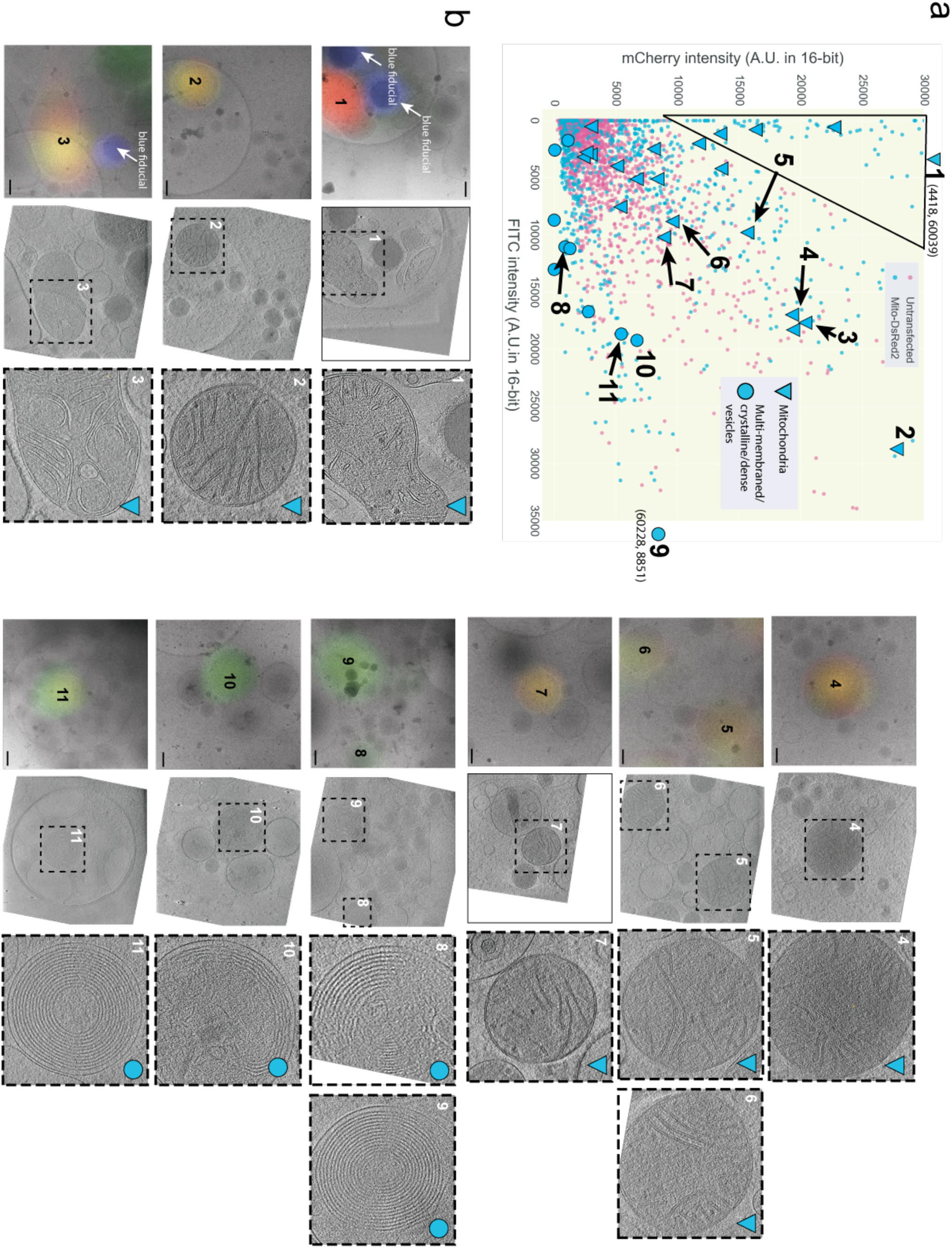
Relative fluorescence intensity can distinguish target fluorescent signal from autofluorescence at 80K. **(a)** Scatter plot of mCherry and FITC channel intensity values of fluorescent puncta in INS-1E cells untransfected (magenta) or transfected with Mito-DsRed2 (blue). [n > X00 for transfected, X00 untransfected.] The black line indicates the area of the scatter plot with no puncta in untransfected INS-1E cells and is described in the text. Larger symbols denote structures observed by cryo-CLEM of selected puncta in transfected cells, corresponding to mitochondria (triangles) or multi-membraned structures, dense vesicles, crystalline structures or structures with a combination of these features (circles). **(b)** Examples of structures correlated to fluorescent puncta. Left panels show epi-fluorescence images overlaid on high-magnification cryo-EM projection images. Middle panels show tomographic slices of the same areas. Right panels show zoomed-in views of the boxed areas in the middle panels. Numbers indicate corresponding locations. Scale bars = 200 nm.

### Cryo-CLEM of INS-1E cells transfected with chromogranin A-GFP

Having established a technique to distinguish signal from autofluorescence, we returned to INS-1E cells expressing CgA-GFP. Once again, we imaged both untransfected and transfected cells by cryo-LM, recorded their intensity values in the green (FITC) and red (mCherry) channels, and plotted these values in two dimensions (Figure 5a). As expected, we observed a broad overlap region in the center of the plot with puncta from both the transfected (tagged) and untransfected (untagged) cells. We also observed a region in the upper left corner of the plot devoid of puncta from untransfected cells which very likely contained CgA-GFP-specific signal.

Twenty-seven puncta were then imaged by ECT, including 15 in the upper left corner of the plot and 12 in the ambiguous overlap region. Cryotomographic slices through ten examples are shown in Figure 5b (see also Supplementary Movie 1). Among the puncta from the upper left corner of the plot (very likely to contain CgA-GFP), we observed vesicles with a dense aggregate core (#’s 4 and 6 for example), vesicles with a dense granular core (#5 for example), vesicles with a dense aggregate or granular core and internal smaller vesicles (#’s 1 and 3 for example), and clusters of dense aggregated material partially surrounded by membrane fragments (#’s 2 and 7). Among the puncta in the ambiguous overlap region we saw two vesicles with crystalline cores (#’s 8 and 10) and a cluster of very dense aggregated material (# 9).

**Figure 5.**
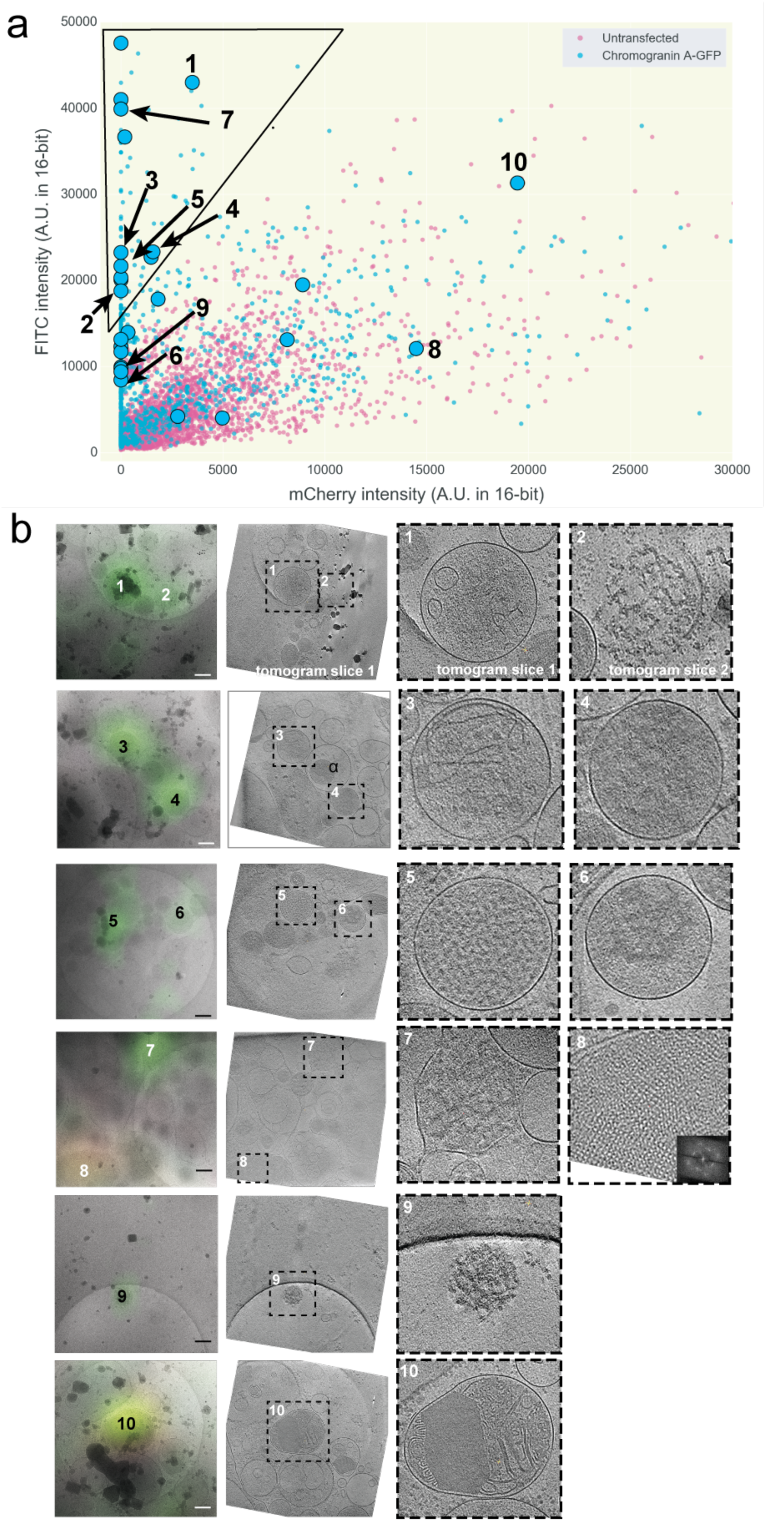
Application of method enables cryo-CLEM of CgA-GFP in INS-1E cells. **(a)** Scatter plot of FITC and mCherry channel intensity values of fluorescent puncta in INS-1E cells untransfected (magenta) or transfected with CgA-GFP (blue). [n > X00 for transfected, X00 untransfected.] The black line indicates the area of the scatter plot with no puncta in untransfected INS-1E cells and is described in the text. Larger symbols denote structures observed by cryo-CLEM of selected puncta in transfected cells (circles). **(b)** Examples of structures correlated to fluorescent puncta. Left panels show epi-fluorescence images overlaid on high-magnification cryo-EM projection images. Middle panels show tomographic slices of the same areas. Right panels show zoomed-in views of the boxed areas in the middle panels. Numbers indicate corresponding locations. Scale bars = 200 nm.

## Discussion

Here we report the discovery that many mammalian cell lines exhibit strong punctate autofluorescence at ∼80K and an approach to distinguish fluorescent protein tags from this autofluorescence.

Cellular autofluorescence at room temperature is known to arise from multiple sources including (1)biomolecules such as amino acids containing aromatic rings, (2) the three-ring system of flavins (producing green spectra) (Benson, Meyer et al. 1979, Jackson, Snyder et al. 2004), (3) the reduced form of pyridine nucleotides (NAD(P)H, producing blue/green spectra) (Chance and Thorell 1959, Galeotti, Van Rossum et al. 1970), (4) lipid pigments (orange/yellow spectra) (Dayan and Wolman 1993, Billinton and Knight 2001), (5) porphyrins (red spectra), and (6) chlorophyll (Sheen, Hwang et al. 1995). Various strategies have been developed to overcome autofluorescence at room temperature including bleaching it away (Viegas, Martins et al. 2007, Kumar, Sandhyamani et al. 2015), quenching it with additives (Cowen, Haven et al. 1985, Schnell, Staines et al. 1999, Billinton and Knight 2001, Kumar, Sandhyamani et al. 2015), or detecting its spectral signature (Steinkamp and Stewart 1986, Van de Lest, Versteeg et al. 1995, Neumann and Gabel 2002, Dickinson, Simbuerger et al. 2003, Gareau, Bargo et al. 2004, Mansfield, Gossage et al. 2005).

Here we found that at ∼80K, autofluorescence is produced by multimembranous structures and what in cell line INS-1E were most likely insulin crystals. The multimembranous structures, which in some cases exhibited up to 14 tightly nested concentric spherical vesicles, resembled MLBs (Hariri, Millane et al. 2000, Lajoie, Guay et al. 2005). Others, bounded by a single outer membrane and containing what looked like partially degraded membranes and other materials are most likely autolysosomes (Klionsky, Eskelinen et al. 2014, Klumperman and Raposo 2014). Earlier work by König et al. also provided evidence that intracellular autofluorescence was more pronounced at 80K than at room temperature, and attributed this to the increase in quantum yield of the fluorophores due to the reduction in thermal relaxation processes at lower temperatures (Konig, Uchugonova et al. 2014).

Since the excitation and emission peaks of autofluorescence overlap with those of commonly used fluorophores such as GFP, YFP and mCherry, filter cubes will not adequately discriminate autofluorescence from signal at 80K. Instead, we recommend first characterizing the autofluorescence by recording images of wild-type (untransfected/unlabelled) cells in two channels, one corresponding to the color of a fluorescent protein tag of interest and the other broadly separated, and then plotting the fluorescence intensities in the two channels in a two-dimensional plot. Next, we recommend imaging a random subset of these puncta by ECT to determine the structures of the sources of autofluorescence in the cell type being studied. Cells labelled with the fluorescent protein tag should then be imaged by cryo-LM in both channels and their fluorescence intensities plotted as before. Puncta exhibiting strong fluorescence in the channel of interest and lying outside the scatter plot region containing puncta from unlabelled cells are very likely to contain the fluorescent protein tag, and not be due simply to autofluorescence. This method requires no specialized equipment and is compatible with a standard cryo-CLEM workflow.

### Cryo-CLEM/ECT of the secretory pathway of INS-1E cells

To demonstrate the utility of the method, we used cryo-CLEM to identify objects containing CgA inside pancreatic cells. These secretory cells are characterized by the ability to rapidly release large amounts of proteins through specialized trans-Golgi network (TGN)-derived vesicles which by traditional EM of stained, plastic-embedded sections display dense granular cores (Kelly 1985, Burgess and Kelly 1987). The dense nature of these cores is known to derive from chromogranins and secretogranins which aggregate in the TGN as their environment acidifies and becomes calcium-rich prior to vesicle formation, and is also related to increased zinc concentrations which are thought to induce insulin crystallization (Gerdes, Rosa et al. 1989, Gorr, Shioi et al. 1989, Chanat and Huttner 1991, Yoo and Albanesi 1991, Videen, Mezger et al. 1992, Taupenot, Harper et al. 2005, Lemaire, Ravier et al. 2009). Aggregation of granin proteins is thought to facilitate cargo sorting (Burgess and Kelly 1987, Carnell and Moore 1994, Taupenot, Harper et al. 2003).

Using our method to identify structures that contained CgA-GFP, we observed not just one class of DCSGs, but a diversity of objects including vesicles with a dense aggregate core, vesicles with a granular core, vesicles with a dense aggregate or granular core and internal smaller vesicles, clusters of dense aggregated material partially surrounded by membrane fragments, and a cluster of very dense aggregated material in the cytoplasm with no membrane fragments in the vicinity. We speculate that the first two classes represent different steps in the secretory pathway, and that the last two classes are the result of vesicle lysis. The vesicles with smaller vesicles inside may be autophagasomes, which are known to degrade insulin as a mechanism to regulate secretory function, though they were not clearly surrounded by two membranes as expected for autophagosomes (Marsh, Soden et al. 2007, Goginashvili, Zhang et al. 2015, Liu, Xue et al. 2015). Compared to the conclusions one might have made based on fluorescence microscopy alone (that all puncta represented DCSGs), cryo-CLEM revealed that some of the puncta were lysed vesicles and probable autophagasomes, which may have formed here simply because of the unnatural levels of CgA expression. Moreover, we also observed vesicles by ECT with dense aggregated cores that were not fluorescent (example denoted by α in Figure 5b; see also Supplementary Movie 1). One possible explanation for this is that the variable pH within DCSGs effects the fluorescence of GFP. Indeed in insulin containing beta cells, granule acidification is a critical step for proper maturation of pro-insulin to the mature form, ultimately leading to crystallization and exocytosis (Orci, Ravazzola et al. 1986, Paroutis, Touret et al. 2004). Again this highlights a caveat to interpreting fluorescence images: some cellular objects of a given type may not fluoresce or even incorporate tagged protein if it is expressed unnaturally. Finally, our images reveal that some vesicles are largely filled by crystals. Because their lattice spacings matched those of insulin crystals, this suggests that insulin can occupy a remarkably large proportion of the vesicle. In any case our cryo-CLEM/ECT data make it clear that the structures of DCSGs are not uniform, but remarkably diverse, and that further characterization is in order.

## Online Methods

### Cell growth and transfection

Rat insulinoma INS-1E (gift of P. Maechler, Université de Genève) and human neuroblastoma BE(2)-M17 cells (CRL-2267; American Type Culture Collection, Manassas, VA) were maintained in a humidified 37°C incubator with 5% CO_2_. INS-1E cells were cultured in RPMI 1640 media with L-glutamine (Life Technologies, Grand Island, NY), supplemented with 5% fetal bovine serum (heat inactivated), 10 mM HEPES, 100 units/mL penicillin, 100 μg/mL streptomycin, 1 mM sodium pyruvate, and 50 μM 2-Mercapto-ethanol. HeLa cells, rhesus macaque fibroblast and primary adipocyte cells were cultured under similar conditions. For cryo-EM and cryo-ET, cells were plated onto fibronectin-coated 200 mesh gold R2/2 London finder Quantifoil grids (Quantifoil Micro Tools GmbH, Jena, Germany) at a density of 2×10^5^ cells/mL. After 48 h incubation, cultures were pre-treated (30 min, 37°C, 5% CO_2_) in Krebs Ringers Bicarbonate HEPES buffer (KRBH: 132.2 mM NaCl, 3.6 mM KCl, 5 mM NaHCO_3_, 0.5 mM NaH_2_PO_4_, 0.5 mM MgCl_2_, 1.5 mM CaCl_2_, and 10 mM HEPES, and 0.1% bovine serum albumin, pH 7.4) supplemented with 2.8 mM glucose before being plunge frozen in liquid ethane/propane mixture using a Vitrobot Mark IV (FEI, Hillsboro, OR) (Iancu, Tivol et al. 2006)˙. For cell transfections, as above INS-1 E cells were plated onto fibronectin-coated 200 mesh gold R2/2 Quantifoil grids at a 2×10^5^ cells/mL density and cultured for 24-48 h (37°C, 5% CO_2_). The cells were then transfected with 2 μg DNA constructs in serum-free RPMI media (5 h, 37°C, 5% CO_2_) using Lipofectamine 2000 (Life Technologies, Carlsbad, CA). Following 16 h incubation in serum-containing RPMI media, cells were washed in KRBH and plunge-frozen for subsequent imaging. Immediately prior to plunge-freezing, 3 μl of a suspension of beads was applied to grids. The bead suspension was made by diluting 500 nm blue (345/435 nm) polystyrene fluorospheres (Phosphorex) with a colloidal solution of 20 nm gold fiducials (Sigma Aldrich) pretreated with bovine serum albumin. The gold served as fiducial markers for tomogram reconstruction while the blue fluorospheres served as landmarks for registering fluorescence light microscopy (FLM) images from different channels as well as EM images. In addition, the blue fluorospheres helped locate target areas in phase contrast light microscopy and low-magnification EM images containing thin ice suitable for high-resolution ECT. Plunge-frozen grids were subsequently loaded into Polara EM cartridges (FEI). EM cartridges containing frozen grids were stored in liquid nitrogen and maintained at ≤−150°C throughout the experiment including cryo-FLM imaging, cryo-EM imaging, storage and transfer.

### Fluorescence imaging and Image processing

The EM cartridges were transferred into a cryo-FLM stage (FEI Cryostage) modified to hold Polara EM cartridges (Nickell, Kofler et al. 2006, Briegel, Chen et al. 2010) and mounted on a Nikon Ti inverted microscope. The grids were imaged using a 60 X extra-long working distance air objective (Nikon CFI S Plan Fluor ELWD 60x NA 0.7 WD 2.62-1.8 mm). Images were recorded using a Neo 5.5 sCMOS camera (Andor Technology, South Windsor, CT) using a real-time deconvolution module in the NIS Elements software (Nikon Instruments Inc., Melville, NY). The pixel size corresponding to the objective lens was ∼108 nm (at the sample level). All fluorescence images (individual channels) were saved in 16-bit grayscale format. CgA-GFP was visualized with a FITC filter. Mito-dsRed2 was visualized with an mCherry filter. Blue fluorospheres were visualized with a DAPI filter. Following FLM imaging, images from different channels were aligned (as described above) using either a module in the NIS Elements software or a Python alignment script written in-house. The 500 nm blue fluorospheres were used to align the DAPI, FITC and YFP channels while autofluorescent puncta were used to align the mCherry channel with the others. The fluorescence channels were aligned to subpixel accuracy. Before the FLM images were further analyzed, background fluorescence in each channel was subtracted from the respective images. This background fluorescence was uniform throughout each image (even outside cellular areas) and likely originates from the grid/ice. Fluorescent puncta were identified in the channel of interest using an in-house python script and their peak fluorescence intensities measured. In addition, intensities of the same pixels in other channels were also recorded. For example, in the case of CgA-GFP dataset, peak intensities of puncta in the FITC channel and their corresponding pixel intensities in the mCherry channel were recorded. Peak intensity values in both channels of both untransfected and transfected cells were plotted on scatter plots.

### Cryo-CLEM and ECT

Grids previously imaged by FLM were subsequently imaged by ECT using an FEI G2 Polara 300kV FEG TEM equipped with an energy filter (slit width 20 eV for higher magnifications; Gatan, Inc.). Images were recorded using a 4k × 4k K2 Summit direct detector (Gatan, Inc.) operating in the electron counting mode. First, areas containing the fluorescent puncta of interest were located in the TEM. Tilt series were then recorded of these areas using UCSF Tomography (Zheng, Keszthelyi et al. 2007) or SerialEM (Mastronarde 2005) software at a magnification of 18,000X. This corresponds to a pixel size of 6 Å (at the specimen level) and was found to be sufficient for this study. Each tilt series was collected from −60° to +60° with an increment of 1° in an automated fashion at 8-10 μm underfocus. The cumulative dose of one tilt-series was between 80 and 200 e^-^ /Å^2^.

Areas of interest were located by TEM in a stepwise manner. First, the grid square/cell of interest on the finder grid was located by using large location markers or other features visible at a low magnification (100 X). Second, a smaller area containing the fluorescent punctum of interest was located by mapping FLM images to intermediate-magnification EM images (typically 3,000X or 1,200X) by eye using various local features within the identified grid square. These features included (1) clusters of 500 nm microspheres that were arranged in a uniquely identifiable pattern,(2)cracks and regularly spaced 2 μm holes in the carbon film and (3) ice contamination. This was done with either UCSF Tomography (Zheng, Keszthelyi et al. 2007) or SerialEM (Mastronarde 2005). With UCSF Tomography, the area of interest first had to be mentally mapped on the low/intermediate magnification EM image using the local features described above before being identified again at 18,000 X magnification by the same features (if available within the field of view). Correlation with SerialEM was more streamlined. FLM images were registered with EM images of the grid square of interest using local features (described above) as control points. These EM images could be either single projection images at a low enough magnification (360X or 1,200X) to contain the area of interest and enough control points to enable tilt-series collection or a montage assembled from higher-magnification (3,000X) EM images. Once the FLM images were registered, areas of interest were marked using the “anchor maps” feature. Using this feature, marked areas could be revisited and tilt-series collected in an automated fashion. Once acquired, tilt-series were aligned and binned four-fold into 1k × 1k arrays before reconstruction into 3D tomograms with the IMOD software package (Kremer, Mastronarde et al. 1996). In addition to the tilt-series, projection images of the location at various magnifications (360X, 1,200X, 3,000X, 9,300X and 18,000X) were saved and used for high-precision post-data collection correlation.

### High-precision post-data collection correlation of FLM images and tomographic slices

RGM fluorescence images were correlated to EM projection images using an in-house image registration script written in Python. This entailed stepwise registration of images recorded at various magnifications, starting with the FLM image, recorded at the lowest magnification (60X), all the way up to the 18,000X EM image. First, the FLM image was registered with a low/intermediate-magnification EM image. The centroid positions of 500 nm blue microspheres were estimated to sub-pixel accuracy and used as control points for this registration. The magnification of the EM image used for this registration was chosen such that there were at least 4 control points available in the field of view. The microspheres were clearly visible at magnifications above 1,200X and less reliably visible at magnifications as low as 360X. Therefore the registration process was more accurate when using >1,200X EM images. The registration parameters (affine transformation) were saved. Successive steps involved similar calculation of affine transformation parameters between EM projection images of various magnifications up to 18,000X. Gold fiducials, surface ice contaminations, and cellular features visible at these magnifications were used as control points for these registration steps. In general, we found that high defocus values at lower magnifications (∼50 μm at 3,000X and ∼1 mm at 1,200X) enhanced the visibility of control points and thus resulted in better registration. Also, having more control points (at least 5) resulted in better registration. Precise registration at lower magnifications is particularly important because of the relatively larger pixel sizes involved. To produce the final registration of the FLM images with the 18,000X projection EM images, the transformations calculated in each of the previous steps were successively applied to the FLM images. The resulting FLM images were overlaid on the 18,000X EM projection images using Adobe Photoshop CC (San Jose, CA), setting the visibility of the upper FLM image layer to linear dodge.

## Acknowledgments

This work was supported by the NIH (grant GM082545 to G.J.J., grant GM082545-6935 to T.J.H., grant K08 DA031241 to Z.F, the Department of Defense (grant PR141292 to Z.F.), and the John F.and Nancy A. Emmerling Fund of The Pittsburgh Foundation (to Z.F.). We thank Dr. Joachim Frank, Dr. Maïté Courel, Dr. Hans Breunig, Robert Grassucci and Stephanie Siegmund for guidance, suggestions and reagents. We thank Dr. Pierre Maechler for generously providing INS-1E cells for our studies.

